# Posterior Periodic Alpha Power as a Neural Marker of Early Biliteracy Skills: The Mediating Role of Rapid Automatized Naming in Chinese-English Bilingual Children

**DOI:** 10.64898/2026.06.10.731265

**Authors:** Tak Kwan Lam, Shuting Huo, Kelvin Fai Hong Lui, Catherine McBride, Urs Maurer

## Abstract

In school-aged children, resting-state posterior alpha activity has been linked to thalamocortical signal coordination and literacy-related cognition. However, findings are mixed, partly because conventional band power measures conflate oscillatory (periodic) activity with the broadband aperiodic (1/f) background. We tested Chinese-English bilingual children in Grades 1 to 5 (N = 121; 72 families; mean age = 8.23 years, SD = 0.81) to examine if posterior periodic alpha power (8–12 Hz), isolated from aperiodic components, predicts foundational biliteracy skills (Chinese and English word reading and dictation). We also tested whether Chinese digit rapid automatized naming (CDRAN) mediates these associations. In regression models that accounted for family clustering and controlled for age, socioeconomic status (SES), and aperiodic components (exponent and offset), higher periodic alpha power uniquely predicted better performance on all four literacy outcomes. In structural equation models that accounted for family clustering, periodic alpha power predicted CDRAN (β = .213, *p* = .011), and CDRAN significantly predicted Chinese word reading (β = .442, *p* < .001), Chinese dictation (β = .346, *p* = .003), English word reading (β = .302, *p* < .001), and English dictation (β = .349, *p* < .001). Indirect effects via CDRAN were significant for all outcomes. These findings suggest that aperiodic-adjusted periodic alpha power is associated with biliteracy variation across Chinese and English and that its association with biliteracy operates in part through rapid serial naming efficiency.

## 1. Introduction

Foundational literacy skills acquired during the primary-school years are a key predictor of children’s later academic success. Reading and spelling require the rapid coordination of visual, attentional, phonological, lexical, and motor processes within a short time window, and this coordination develops alongside the maturation of the brain systems that support learning. Resting-state electroencephalography (EEG) provides a well-tolerated, non-invasive method for capturing millisecond-level brain dynamics in children, and posterior alpha activity is a promising candidate neural marker because of its developmental salience and theoretical links to thalamocortical timing and inhibitory control (Biasiucci et al., 2019; Bollimunta et al., 2011; Hughes & Crunelli, 2005; Klimesch et al., 2007). Yet the literature relating resting-state alpha to literacy has been mixed. The present study addresses three questions that emerge from this literature: whether alpha-literacy associations become clearer when periodic alpha is separated from aperiodic background activity; whether such associations generalize beyond Chinese first-language literacy to Chinese-English biliteracy; and whether rapid automatized naming (RAN) helps explain the relation between posterior alpha and literacy outcomes.

### 1.1 Periodic Alpha Oscillations, Aperiodic Activity, and Literacy

A prominent posterior dominant rhythm is present across childhood, but its peak frequency is typically slower in younger children and shifts upward with age toward the canonical 8-12 Hz alpha range (Freschl et al., 2022; Tröndle et al., 2022). Alpha oscillations are primarily generated and regulated by thalamocortical circuits. Through reciprocal thalamocortical loops, nuclei such as the pulvinar and the lateral geniculate nucleus can synchronize widespread cortical activity, providing a timing and gating mechanism that helps coordinate information flow across distributed networks (Hughes & Crunelli, 2005; Llinás, 1993). Consistent with the inhibition-timing hypothesis, alpha oscillations are thought to suppress task-irrelevant activity while facilitating processing in task-relevant networks, thereby creating a temporal window for efficient cognition (Klimesch et al., 2007; Schoffelen et al., 2024).

These theoretical links make posterior alpha relevant to literacy development, which depends on the rapid coordination of multiple cognitive processes (Norton & Wolf, 2012; Schlaggar & McCandliss, 2007). Stronger resting-state alpha power has been associated with cognitive-linguistic skills relevant to reading, such as vocabulary and working memory, and with reading-related outcomes such as fluency (Harmony et al., 1990; Maguire & Schneider, 2019; Myers et al., 2014; Vogt et al., 1998; Whedon et al., 2020). Several studies have also reported atypical resting-state alpha in children with dyslexia (Babiloni et al., 2012; Xue et al., 2020). At the same time, findings have not been fully consistent, and some studies have implicated other frequency bands or reported weak relations between resting alpha and language-related outcomes (Arns et al., 2007; Fein et al., 1986; Kwok et al., 2019; Schneider et al., 2021).

One plausible reason for these inconsistencies is methodological. Conventional band-power measures treat the EEG spectrum as a unitary signal, even though the spectrum can be decomposed into dissociable periodic and aperiodic components (Donoghue et al., 2020; He, 2014; Smit et al., 2011). Periodic activity reflects narrowband oscillatory peaks, whereas aperiodic activity reflects the broadband 1/f-like background. Developmental work shows that these components follow distinct trajectories when modelled separately, so apparent changes in “alpha power” can partly reflect changes in aperiodic activity rather than changes in rhythmic alpha itself (Hill et al., 2022; Rico-Picó et al., 2023; Tröndle et al., 2022; Wilkinson et al., 2023). The same confound may also apply to individual differences, such that similar total alpha power can arise from different combinations of periodic and aperiodic activity. Separating periodic alpha from the aperiodic background therefore provides a cleaner test of whether oscillatory alpha is specifically associated with literacy-related behaviour.

### 1.2 Biliteracy as a Test of Generalization Across Writing Systems

Most prior work on resting-state alpha and literacy has focused on first-language reading. A Chinese-English bilingual sample offers an opportunity to test whether the association between posterior alpha and literacy generalizes across languages that differ markedly in orthographic structure. During the early school years, children in bilingual contexts develop biliteracy, which is the ability to read and write in two languages, through both language-specific knowledge and shared processing resources. Cross-linguistic accounts of literacy development propose that some abilities supporting literacy, such as processing efficiency, can transfer across languages even when scripts differ substantially (Goodrich & Lonigan, 2017; Verhoeven, 1994).

If posterior periodic alpha indexes shared timing and coordination mechanisms rather than language-specific representations, its association with literacy should not be limited to Chinese. Examining both Chinese and English tests whether the same neural marker relates to literacy in both a first language and a second language, and across logographic and alphabetic writing systems. The present study focused on two core literacy outcomes in both languages—word reading and dictation. Dictation is especially informative because it requires the integration of orthographic knowledge, phonological processing, and working memory, and in Chinese it is also predicted by RAN (Kalindi & Chung, 2018).

### 1.3 Rapid Automatized Naming as a Behavioural Pathway

A further question is whether the alpha-literacy association is partly mediated by rapid automatized naming. RAN performance depends on the efficient coordination of visual scanning, attention, access to overlearned verbal labels, and articulation under time pressure (Wolf et al., 2000). From a thalamocortical timing perspective, stronger and more stable posterior alpha may index more efficient neural timing for such rapid serial processing (Jensen et al., 2012; Lakatos et al., 2008; Samaha & Postle, 2015). This proposal is relevant because RAN is one of the most robust predictors of reading and spelling across languages and writing systems, including Chinese (Chen et al., 2021; Georgiou et al., 2016; Norton & Wolf, 2012; Wolf & Bowers, 1999; X. Yang et al., 2024), and can show cross-language associations in bilingual children (Kishchak et al., 2024).

### 1.4 Current Study and Hypotheses

The present study was designed around three linked aims. First, we examined whether periodic alpha power predicts literacy more clearly when it is separated from aperiodic components. As a supplementary comparison, we also tested whether the same associations would be detectable using conventional total alpha power. Second, we tested whether the association generalizes across Chinese and English word reading and dictation in Chinese-English bilingual children. Third, we examined whether CDRAN mediates the relation between periodic alpha power and biliteracy outcomes. All analyses controlled for age and SES, and all models accounted for the clustering of twins within families.

We expected higher periodic alpha power to be associated with better performance on Chinese word reading (CWR), Chinese dictation (CDICT), English word reading (EWR), and English dictation (EDICT) above and beyond the aperiodic exponent and offset. We further expected this pattern to be present in both Chinese and English, consistent with a shared cross-language contribution of posterior alpha to literacy. Finally, we expected higher periodic alpha power to predict better CDRAN performance, which in turn would predict stronger literacy outcomes in both languages.

## 2. Methods

### 2.1 Participants

Children were recruited from a large-scale longitudinal twin study in Hong Kong (Chung et al., 2023; Lo et al., 2019). Resting-state EEG data from this cohort have been reported previously (Lui et al., 2021). We initially selected a total of 202 children from 106 families with usable EEG recordings from Grades 1 to 5 (*M* age = 8.09 years, *SD* = 0.98, 60 boys). After excluding participants with missing behavioural data, and after EEG preprocessing procedure, the final analytic sample comprised 121 children from 72 families remained (*M* age = 8.23 years, *SD* = 0.81, 47 boys). All participants were native Cantonese speakers with no reported history of neurodevelopmental diagnoses (including developmental dyslexia, intellectual disability, autism spectrum disorder, or attention-deficit/hyperactivity disorder) and had normal or corrected-to-normal vision. Parents provided informed consent. The study protocol was approved by the University Ethics Committee.

### 2.2 Procedure and EEG Recording

Children completed two sessions with behavioural assessment and resting-state EEG recording, held on average within 2 months of each other. Behavioural testing took place at home or at the child’s primary school. EEG was recorded in a sound-attenuated laboratory using a HydroCel GSN 128-channel system (Electrical Geodesics Inc., Eugene, OR, USA) with Net Station software.

During EEG acquisition, children completed a 3-minute eyes-open resting-state recording. They were seated approximately 80 cm from the monitor and instructed to remain as still as possible while fixating a central cross for the duration of the recording.

### 2.3 Early Biliteracy Skills

#### 2.3.1 Chinese Word Reading (CWR)

Children were assessed by an adapted version of The Hong Kong Test of Specific Learning Difficulties in Reading and Writing (Ho et al., 2000), which includes 150 two-character words. The items were arranged in order of increasing difficulty. The task would stop when the child failed to read 15 consecutive items. The total number of words they read correctly served as the indicator of their word reading performance (maximum = 150). Split-half reliability was excellent (Guttman’s λ4 = 0.96), and Cronbach’s alpha was 0.93.

#### 2.3.2 Chinese Dictation (CDICT)

A test from Ho et al., (2000) was used for the CDICT task, but only with 20 two-character Chinese words. All the words to be spelled on this task were representative of beginning words taught in Hong Kong, with the order arranged by increasing difficulty. Testing was stopped when children missed five consecutive words in a row. One mark was given for each correctly written word, so the maximum score was 40. Split-half reliability was good (Guttman’s λ4 = 0.86), and Cronbach’s alpha was 0.87.

#### 2.3.3 English Word Reading (EWR)

Children were assessed by the adapted version of reading list from (Tong & McBride-Chang, 2010). All words were selected from Hong Kong Chinese children’s English textbooks that were frequently used in kindergarten and primary school. Ten easy items were deleted from the list and thus the test consisted of 50 English words in total. The test items were arranged in 5 levels according to reading difficulty. Testing stopped when children had 4 consecutive wrong answers within one level. The maximum score of this task was 50. Split-half reliability was excellent (Guttman’s λ4 = 0.95), and Cronbach’s alpha was 0.90.

#### 2.3.4 English Dictation (EDICT)

A test from Lo et al. (2018) was used for the EDICT task. A total of 12 English compound words (e.g., nighttime) were presented to children with an mp3 player and increasing difficulty. One point was awarded for each correctly spelled morpheme (max score = 24). Split-half reliability was good (Guttman’s λ4 = 0.85), and Cronbach’s alpha was 0.85.

### 2.4 Chinese Digit Rapid Automatized Naming (CDRAN)

The rapid automatized naming task was adapted from the rapid automatized naming task by Denckla and Rudel (1976). The task consisted of eight rows of five digits [e.g., 2, 4, 6, 7, and 9]. These digits were arranged in different orders for each row. Children were asked to name the digits in Cantonese as quickly and accurately as possible. Two trials were completed and the average time in seconds was recorded. First, we calculated the mean time (in seconds) to name one digit. Then, to find the naming speed per minute, we divided 60 seconds by this mean time to get the CDRAN score. Test-retest reliability across the two trials was excellent, ICC (2,1) = 0.87.

### 2.5 EEG Preprocessing and Parameterization

EEG was recorded at 500 Hz with Cz electrode as the online reference and a 0.1 Hz high-pass filter. Electrode impedances were kept below 50 kΩ. We followed the preprocessing pipeline reported by Lui et al. (2021), implemented in EEGLAB v14.1.2 (Delorme & Makeig, 2004). The first and the last 10 seconds of the continuous EEG data were removed to exclude movement-related artifacts. Data were downsampled to 100 Hz to facilitate independent component analyses (ICA). ICA was performed using the CUDAICA Infomax optimization algorithm (Raimondo et al., 2012), and ocular components were identified and removed using ADJUST (Mognon et al., 2011). The data were then filtered with a 40 Hz low-pass filter and then segmented into adjacent 4-second epochs. Epochs with absolute voltage values exceeding 100 μV at any electrodes were removed. We required a minimum of 15 artifact-free 4-s epochs (≥ 60 s) for inclusion, consistent with prior pediatric resting-state EEG work that excludes participants with < 1 minute of usable data after preprocessing (Lui et al., 2021; Y. Yang et al., 2025). All participants retained at least fifteen artifact-free epochs (≥ 60 s; *M* epochs = 31.38, *SD* = 8.86). Finally, data were re-referenced to the average reference.

Power spectra were computed for each epoch and channel using the fast Fourier transform, and spectra were averaged across epochs. We parameterized the power spectrum in the 2–25 Hz range using the Fitting Oscillations and One-Over-F (FOOOF) algorithm to separate periodic alpha oscillation from aperiodic components in power density spectrum (Donoghue et al., 2020), thereby separating periodic peaks from the aperiodic (1/f) background. Following Huo et al. (2024), we constrained the maximum number of fitted peaks to four and set the minimum peak width to 2 Hz. Periodic alpha power was defined as the amplitude of the strongest aperiodic-adjusted oscillatory peak within 8–12 Hz.

To focus on the canonical posterior alpha generators, we quantified periodic alpha power from an occipito-parietal region of interest. After fitting the spectral parameterization model for all 128 channels, we visualized the sample-average topography of the aperiodic-adjusted alpha peak amplitude (Figure 1a), which showed a posterior maximum consistent with prior resting-state alpha work (Liu et al., 2012; Patel et al., 1999). We then identified the three channels with the highest mean periodic alpha power across participants (E62, E79, E87), and averaged of their periodic alpha peak amplitudes for each child. This three-channel average served as the periodic alpha predictor in all analyses (Figure 1b).

**Figure 1.**
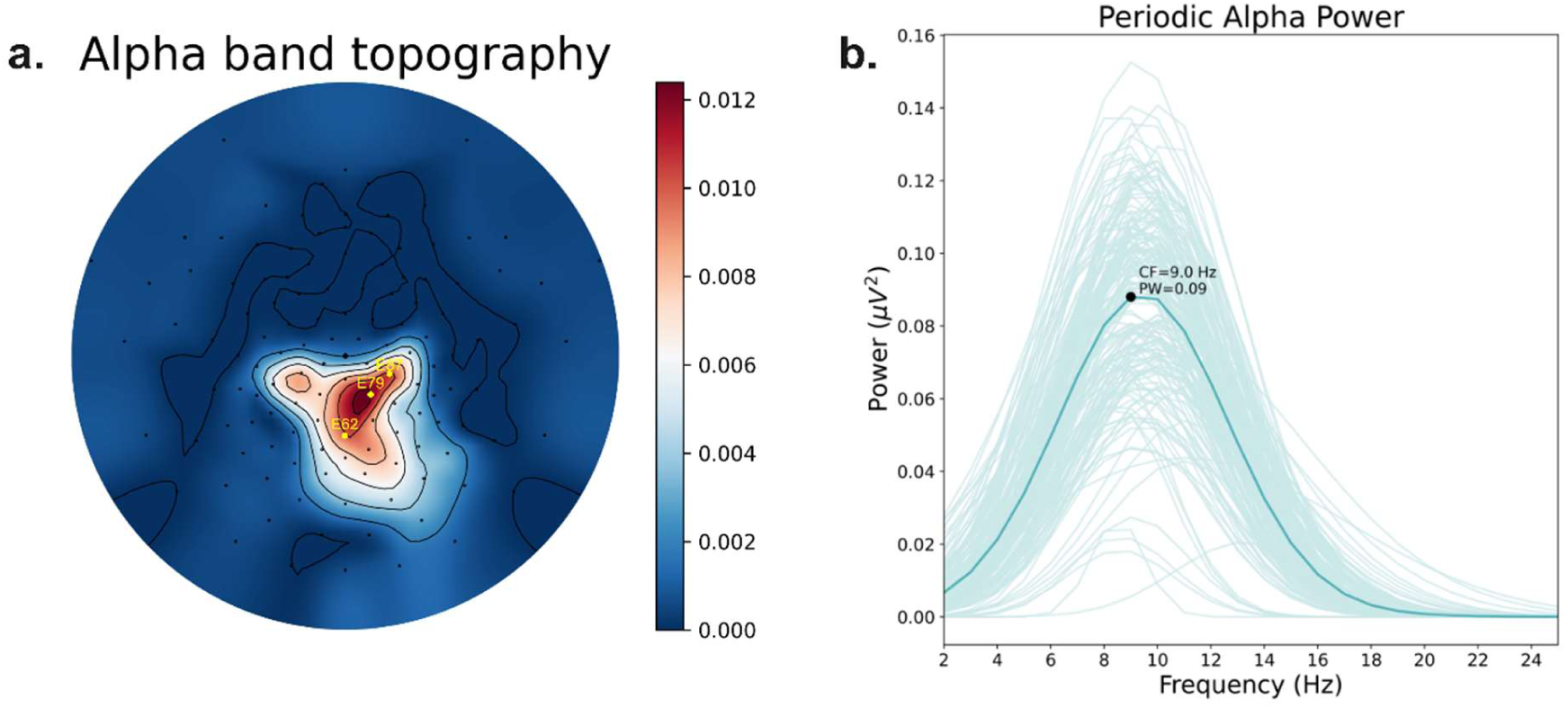
The Topography of the Periodic Alpha Band Peak Power and the Periodic Power Curve of the Children. *Note.* Panel (a) shows the sample-averaged topographic map of the aperiodic-adjusted peak periodic alpha power, averaged across all participants. The three most dominant channels (E62, E79, E87) used for analysis are marked in yellow. Panel (b) shows periodic power spectra from these channels. The dark blue line represents the mean power across children; lighter blue lines represent individual children’s power curves., and the black marker indicates the sample-average peak.

### 2.6 Data Analysis

Given the nested structure of our data (twins within families), we accounted for non-independence using multilevel models (random intercepts for family) and cluster-robust inference where appropriate.

#### 2.6.1 Analytical Strategy for Hypothesis 1 (H1): Regression Models

To test H1, we estimated a family-clustered multivariate regression model in which Chinese word reading (CWR), Chinese dictation (CDICT), English word reading (EWR), and English dictation (EDICT) were regressed simultaneously on periodic alpha power (PW), aperiodic exponent, aperiodic offset, age, and SES, with residual covariances among the literacy outcomes freely estimated. Family clustering was accounted for in the estimation. This model allowed us to test whether oscillatory alpha uniquely predicted literacy after accounting for the aperiodic parameters and shared variance among outcomes. Figure 2 provides covariate-adjusted visualizations of the periodic and aperiodic effects, and Supplementary Table 1 reports the full parameter estimates. As a supplementary comparison, we also fit parallel adjusted models using conventional total alpha power instead of the parameterized predictors; these comparison results are summarized in Table 2.

#### 2.6.2 Structural Equation Model for Hypothesis 2 (H2)

To test H2, we estimated a structural equation model in lavaan v0.6-18 (Rosseel, 2012) with direct paths from periodic alpha power to each literacy outcome and indirect paths through CDRAN. Age and SES were included as covariates on CDRAN and all outcomes. Because the data were clustered by family, the substantive mediation structure was specified at the between-family level, whereas the within-family level captured residual variances and covariances only. Thus, all paths among periodic alpha power, CDRAN, and the literacy outcomes were modelled at the between-family level.

Indirect and total effects were estimated using 5,000 bias-corrected and accelerated (BCa) bootstrap resamples. Statistical significance was evaluated using bootstrap p-values and 95% confidence intervals. Model fit was evaluated using χ²/df, the comparative fit index (CFI), the Tucker-Lewis index (TLI), the root mean square error of approximation (RMSEA), and the standardized root mean square residual (SRMR) (Hu & Bentler, 1999; Kline, 2016).

## 3. Results

### 3.1 Association Between Early biliteracy development, Periodic, and Aperiodic Components in Power Density Spectra

Descriptive statistics and the partial bivariate correlations (controlling for age and socioeconomic status) among all study variables are shown in Table 1. Periodic alpha power (PW) showed significant positive correlations with all four literacy measures: Chinese word reading (CWR: β = .36, *p* = .018), Chinese dictation (CDICT: β = .21, *p* = .015), English word reading (EWR: β = .14, *p* = .037), English dictation (EDICT: β = .17, *p* = .015), and with Chinese digit rapid automatized naming (CDRAN: β = .21, *p* = .009). In contrast, the aperiodic components showed limited significant associations with biliteracy measures. Only the aperiodic offset was significantly correlated with CWR (β = -.16, *p* = .037). The aperiodic exponent was not significantly correlated with any biliteracy variables (all *p* > .05).

**Table 1.**
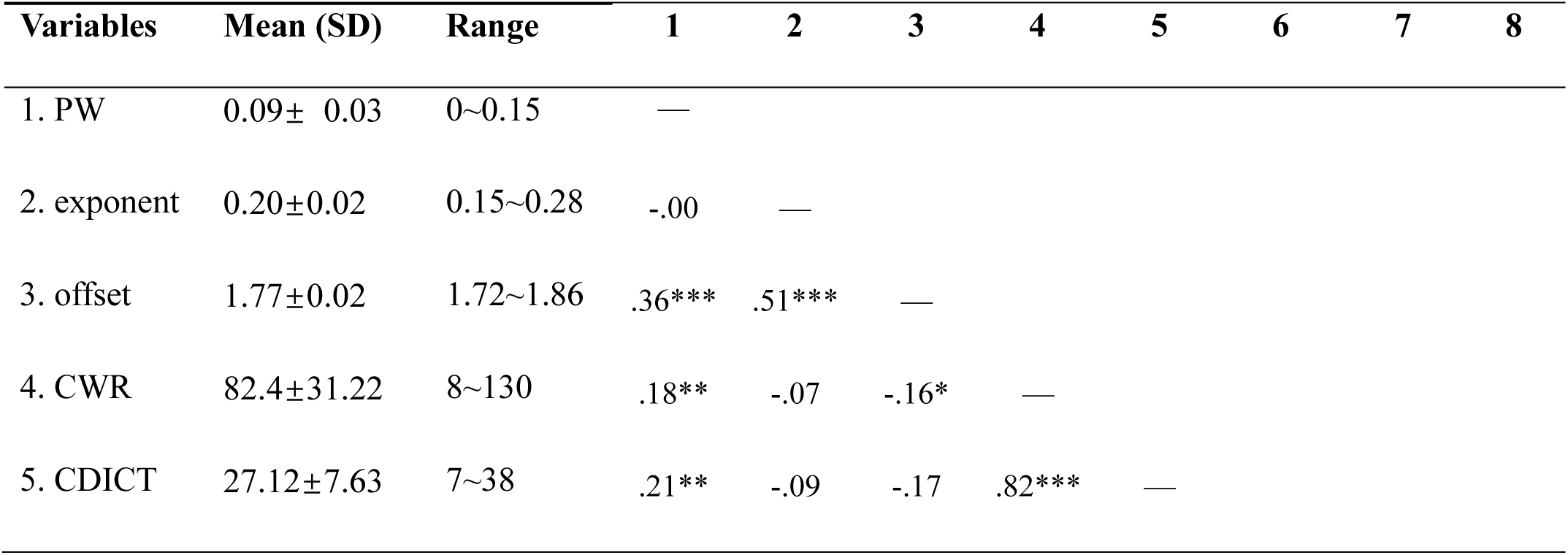
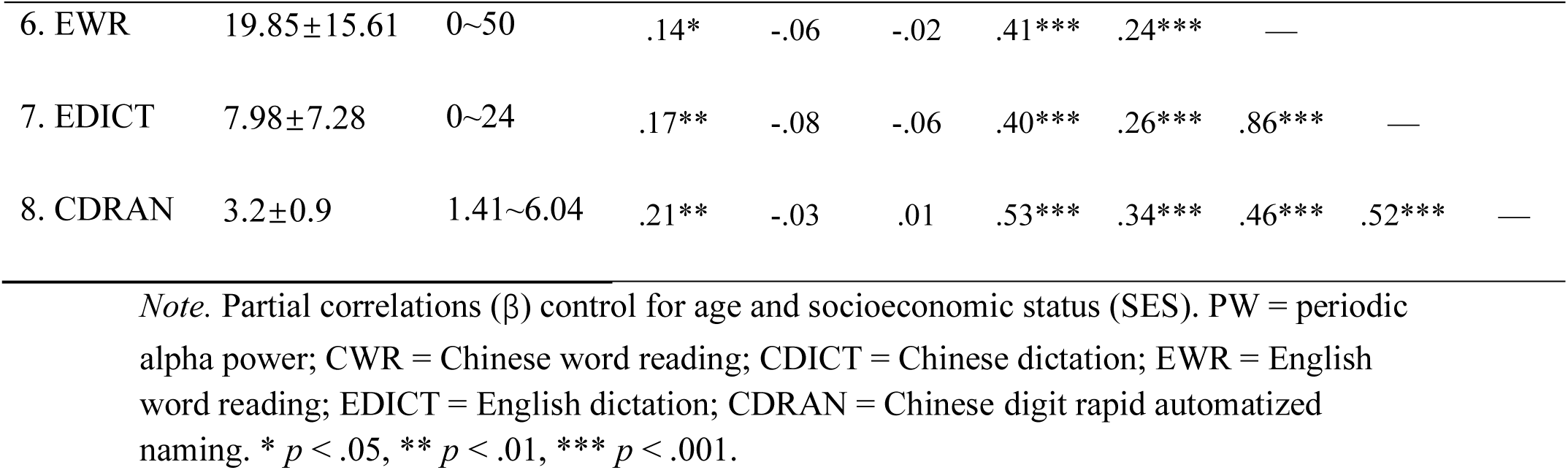
Descriptive Statistics and Bivariate Correlations for Neural and Biliteracy Variables.

### 3.2 Dissociable Contributions of Periodic Alpha, Aperiodic Offset, and Total Alpha Power

To test whether separating periodic and aperiodic components clarified alpha–literacy associations, we first estimated the family-clustered multivariate model reported in Supplementary Table 1 and visualized the adjusted associations in Figure 2. Higher periodic alpha power (PW) was a significant unique predictor of all four biliteracy skills (Figure 2a; Supplementary Table 1): Chinese word reading (CWR: β = .244, *p* < .001), Chinese dictation (CDICT: β = .313, *p* < .001), English word reading (EWR: β = .138, *p* = .028), and English Dictation (EDICT: β = .179, *p* = .011). The aperiodic component exponent did not significantly predict any biliteracy outcome. The aperiodic offset was a significant negative predictor for CWR (β = -.246, *p* = .010) and CDICT (β = -.284, *p* = .001), but not with the English outcomes (Figure 2b; Supplementary Table 1). Thus, the periodic and aperiodic results were dissociated both in direction and in their pattern of generalization across outcomes. Importantly, when we repeated the analysis using conventional total alpha power, none of the literacy outcomes showed a significant association (Table 2). This comparison indicates that the positive periodic alpha association was not detectable when periodic and aperiodic components were not separated. Age significantly predicted all outcomes, and SES significantly predicted English literacy outcomes (Supplementary Table 1).

**Figure 2.**
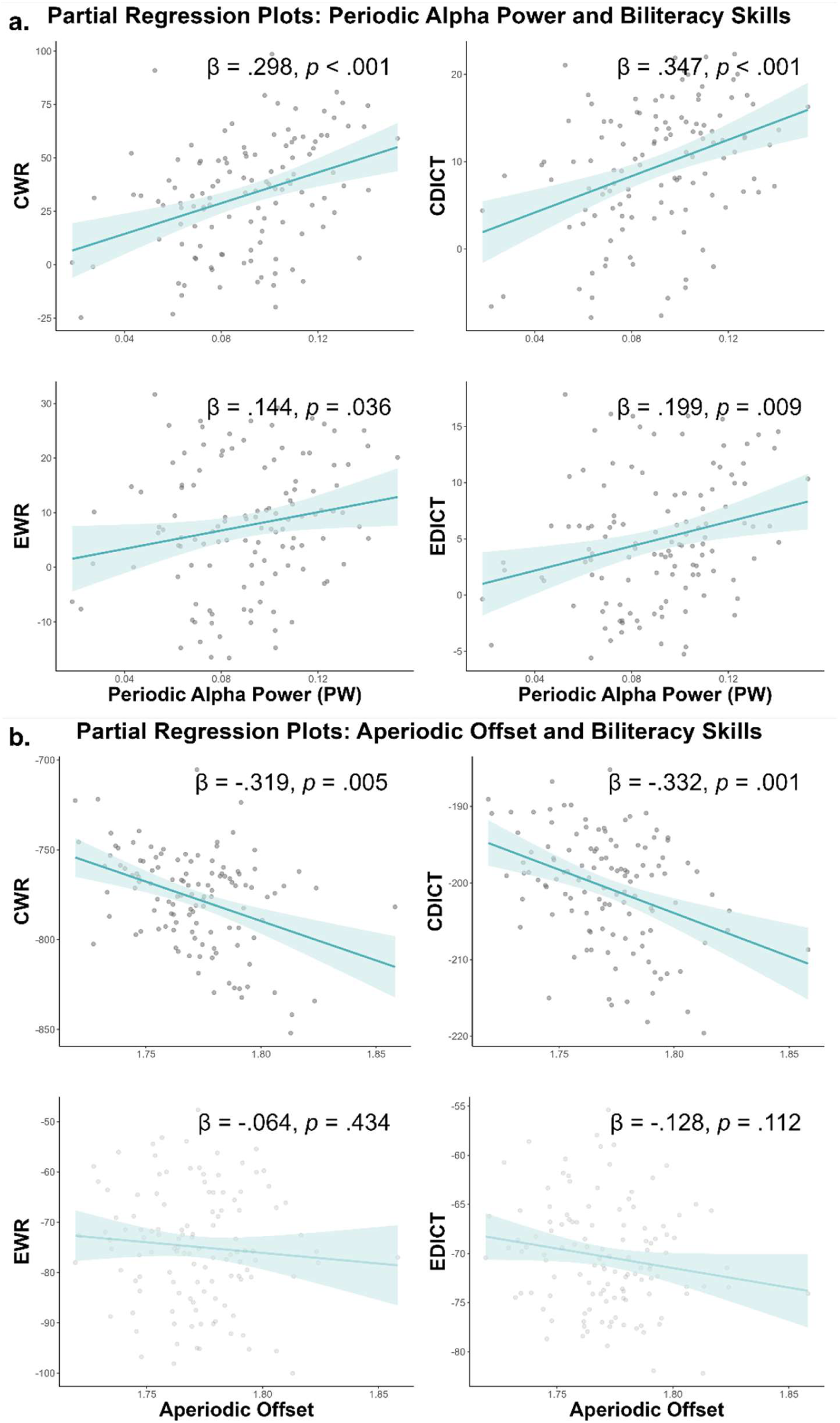
Partial Associations of Periodic Alpha Power and Aperiodic Offset with Biliteracy Outcomes. *Note.* Panel (a) shows covariate-adjusted partial regression plots for periodic alpha power (PW) predicting Chinese word reading (CWR), Chinese dictation (CDICT), English word reading (EWR), and English dictation (EDICT). Panel (b) presents the corresponding plots for aperiodic offset. Each association is adjusted for age, SES, and the remaining spectral parameters. Aperiodic exponent is not shown because it was not a significant predictor of any literacy outcome. The plotted coefficients are shown for visualization; inferential statistics for the family-clustered model are reported in Supplementary Table 1. Darker lines indicate statistically significant associations, whereas lighter lines indicate non-significant associations.

**Table 2.**
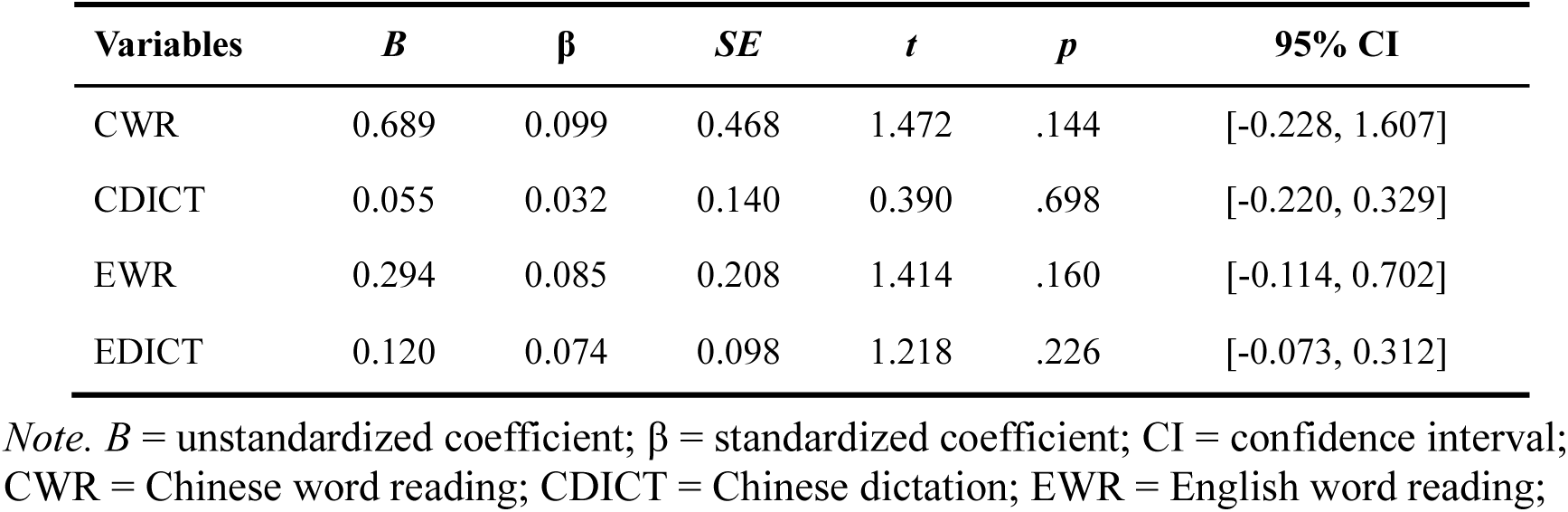
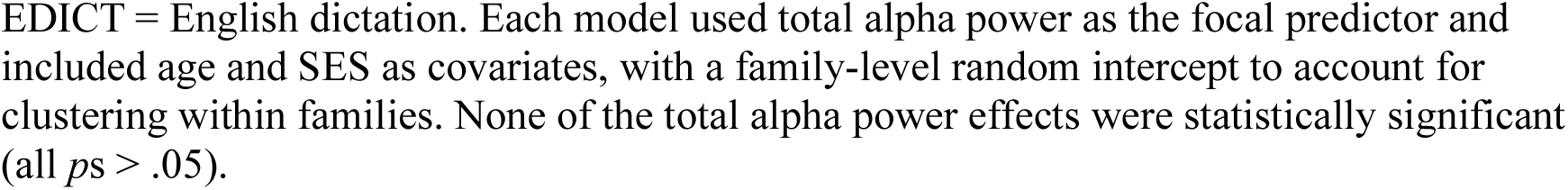
Supplementary Comparison Models Using Conventional Total Alpha Power to Predict Biliteracy Outcomes.

| Variables | *B* | $\beta$ | *SE* | *t* | *p* | 95% CI |
| --- | --- | --- | --- | --- | --- | --- |
| CWR | 0.689 | 0.099 | 0.468 | 1.472 | .144 | [-0.228, 1.607] |
| CDICT | 0.055 | 0.032 | 0.140 | 0.390 | .698 | [-0.220, 0.329] |
| EWR | 0.294 | 0.085 | 0.208 | 1.414 | .160 | [-0.114, 0.702] |
| EDICT | 0.120 | 0.074 | 0.098 | 1.218 | .226 | [-0.073, 0.312] |

*Note.* *B* = unstandardized coefficient; $\beta$ = standardized coefficient; CI = confidence interval; CWR = Chinese word reading; CDICT = Chinese dictation; EWR = English word reading;
EDICT = English dictation. Each model used total alpha power as the focal predictor and included age and SES as covariates, with a family-level random intercept to account for clustering within families. None of the total alpha power effects were statistically significant (all $ps > .05$ ).

### 3.3 Mediation of Periodic Alpha Power through Rapid Automatized Naming

To investigate H2, we estimated a structural equation model that accounted for family clustering and included direct paths from periodic alpha power to each biliteracy outcome and indirect paths through rapid automatized naming (CDRAN), while controlling for age and SES. Model fit was excellent (χ² = 0.480, df = 2, *p* = .787; CFI = 1.000; TLI = 1.059; RMESA = 0.000; SRMR = 0.020). Periodic alpha power significantly predicted CDRAN (β = .213, *p* = .011). CDRAN, in turn, significantly predicted all biliteracy skills: CWR (β = .442, *p* < .001), CDICT (β = .346, *p* = .003), EWR (β = .302, *p* < .001), and EDICT (β = .349, *p* < .001).

The SEM revealed significant indirect effects of periodic alpha power on biliteracy skills through CDRAN for Chinese word reading (*B* = 106.158, β = .094, *p* = .014, 95% CI [24.215, 192.549]), Chinese dictation (*B* = 20.251, β = .074, *p* = .022, 95% CI [4.514, 38.235]), English word reading (*B* = 19.563, β = .074, *p* = .030, 95% CI [7.873, 71.265]), and English dictation (*B* = 36.255, β =.064, *p* = .032, 95% CI [4.113, 38.072]). Significant total effects of periodic alpha power were also observed for all biliteracy skills (CWR: *B* = 204.627, β = .182, *p* = .009, 95% CI [48.489, 360.541]; CDICT: *B* = 58.424, β = .213, *p* = .007, 95% CI [14.783, 106.988]; EWR: *B* = 44.621, β = .170, *p* = .011, 95% CI [9.643, 144.259]; EDICT: *B* = 80.510, β = .143, *p* = .028, 95% CI [10.326, 74.982]). Direct effects from periodic alpha power to biliteracy skills became non-significant for CWR (β = .087, *p* = .177), and EWR (β = .079, *p* = .186), consistent with substantial mediation for word reading. Direct effects were also non-significant for dictation (CDICT: β = .139, *p* = .064; EDICT: β = .095, *p* = .118), suggesting full mediation for both reading and spelling (Figure 2). The model explained substantial variance in the outcome variables: rapid automatized naming (R^2^ = .248), Chinese word reading (R^2^ = .477), Chinese dictation (R^2^ = .236), English word reading (R^2^ = .534), and English dictation (R^2^ = .545).

**Figure 2.**
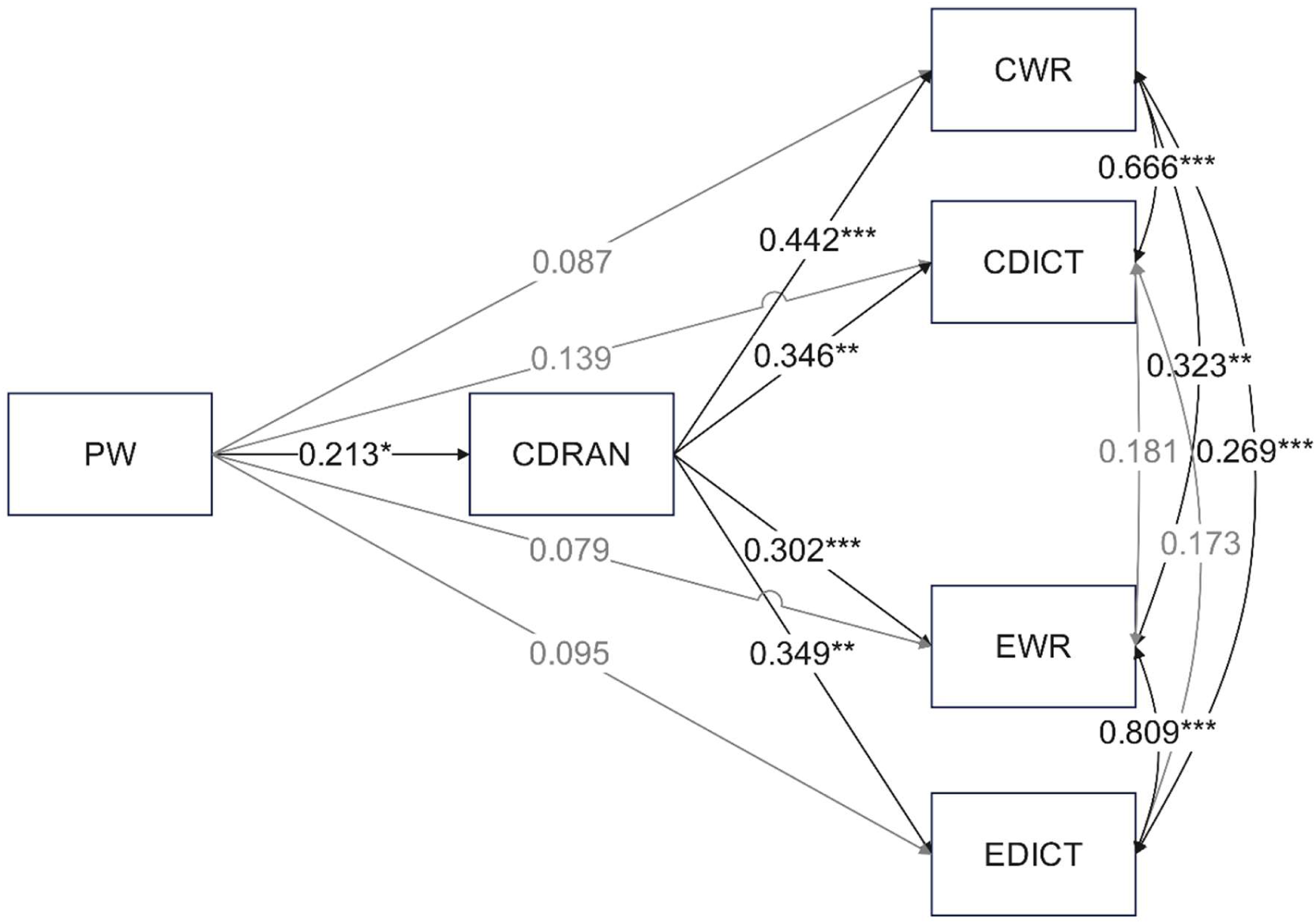
Standardized Structural Equation Model Predicting Biliteracy Abilities. *Note*. Asterisks indicate significance level: *, *p* < .05, **, *p* < .01, ***, *p* < .001. PW = periodic alpha power; CWR = Chinese word reading; CDICT = Chinese dictation; EWR = English word reading; EDICT = English dictation; CDRAN = Chinese digit rapid automatized naming. Gray lines represent non-significant paths (*p* ≥ .05).

## 4. Discussion

The study examined whether posterior resting-state periodic alpha power, isolated from the aperiodic (1/f) background via spectral parameterization, is associated with foundational biliteracy skills in Chinese–English bilingual children, and whether this association is partly accounted for by rapid automatized naming. Three points stand out. First, separating periodic and aperiodic components revealed a clear dissociation: periodic alpha showed positive associations with literacy, whereas aperiodic offset showed negative associations with the Chinese literacy measures. Second, this pattern was observed for both Chinese and English word reading and dictation for periodic alpha, but the offset effect was more selective. Third, conventional total alpha power was not significantly related to any literacy outcome once the spectrum was treated as a single undifferentiated band. Fourth, CDRAN significantly mediated the association between periodic alpha and all four literacy outcomes. Together, these findings support the view that posterior periodic alpha indexes neural processes that are relevant to early literacy and that part of this relation operates through rapid serial naming efficiency.

### 4.1 Periodic Alpha Power and Foundational Biliteracy Skills Across Chinese and English

Our first goal was to clarify the mixed resting-state alpha literature by separating oscillatory alpha from the aperiodic background. Two aspects of the result deserve emphasis. First, higher aperiodic-adjusted periodic alpha power was associated with better Chinese and English word reading and dictation, whereas higher aperiodic offset was associated with poorer Chinese word reading and dictation (Figure 2; Supplementary Table 1). This opposite pattern is theoretically important because it shows that periodic and aperiodic features do not simply provide redundant versions of the same neural signal.

Second, when these components were not separated, conventional total alpha power was not significantly associated with any literacy outcome (Table 2). This null comparison helps explain why the resting-state alpha literature has often been mixed: if a positive periodic association and a negative aperiodic association are blended into a single band-power estimate, the effect can be attenuated or obscured. In this sense, modelling periodic and aperiodic components separately does not merely refine interpretation; it changes whether an alpha–literacy association is detectable at all.

### 4.2 The Association Generalizes Across Chinese and English Literacy

Our second goal was to determine whether the alpha-literacy association extends beyond Chinese first-language literacy to biliteracy across Chinese and English. This was important because Chinese and English differ sharply in orthographic structure. Word reading requires rapid access to orthographic and phonological representations, whereas dictation additionally places demand on holding the item in mind, retrieving the correct spelling, and producing it accurately. yet both rely on efficient coordination of shared cognitive processes such as attention, processing speed, and access to stored orthographic and phonological representations. The fact that periodic alpha predicted both Chinese and English, and for both reading and dictation, is also important. Finding the same general pattern across two writing systems and two types of literacy outcome suggests that posterior periodic alpha is not tied to one narrow skill. Instead, posterior periodic alpha may index shared timing and coordination mechanisms that support literacy performance across languages.

This cross-language pattern should nevertheless be interpreted cautiously. The presence of associations in both Chinese and English does not imply that the same processes contribute equally to both languages. English is a second language for the children in this sample, and the strength of the relation may also depend on exposure, proficiency, and the additional control demands of second-language processing (Abutalebi, 2008; Ye & Zhou, 2009). The present findings therefore support generalization across languages without implying that the underlying developmental pathways are identical.

### 4.3 Rapid Automatized Naming as a Behavioural Pathway

Our third aim was to test whether rapid automatized naming provides a behavioural pathway linking periodic alpha to literacy. The mediation results support this interpretation. Periodic alpha power predicted better CDRAN performance, and CDRAN, in turn, predicted all four literacy outcomes. Because RAN requires children to coordinate visual scanning, sustained attention, access to overlearned verbal labels, and articulation under time pressure, it is a plausible behavioural link between resting neural timing and literacy performance (Wolf et al., 2000). In this sense, the findings are consistent with the idea that stronger posterior alpha reflects more efficient timing or gating processes that support rapid serial access to overlearned symbols.

This interpretation is compatible with prior work linking alpha dynamics to naming speed. Higher prestimulus alpha power in motor regions has been associated with faster picture naming latencies (Jongman et al., 2020), and the broader RAN literature shows that naming speed is a robust predictor of reading and spelling across writing systems (Chen et al., 2021; Georgiou et al., 2016; Kishchak et al., 2024). At the same time, the present study is cross-sectional and based on resting-state EEG. The indirect effects are therefore statistically consistent with mediation, but they should not be interpreted as proof of causal developmental pathways. Other mechanisms, such as orthographic knowledge, phonological processing, and morphological awareness, may also contribute, especially for dictation.

### 4.4 Aperiodic Components and Their Role in Biliteracy Development

A further contribution of the study is that it considered aperiodic parameters alongside periodic alpha rather than treating alpha-band power as a single undifferentiated measure. This mattered empirically. After periodic alpha, age, and SES were controlled, higher aperiodic offset predicted poorer Chinese word reading and dictation, whereas the exponent did not show reliable relations to the literacy outcomes. The offset effect was therefore more selective than the periodic alpha effect and was confined to the Chinese measures. One possible interpretation is that elevated broadband activity reflects a less efficient or noisier baseline neural state (Deodato & Melcher, 2024; Krystecka et al., 2024). If so, a higher offset may be especially unhelpful for Chinese literacy tasks, which place heavy demands on fine-grained visual-orthographic analysis and stable retrieval of character forms. This interpretation should remain cautious, however, because the functional meaning of offset is still less well established than that of periodic alpha. Figure 2 makes the dissociation between periodic alpha and offset visually clear, and the null total-alpha comparison in Table 2 shows its practical consequence: once the spectrum is reduced to a single band-power estimate, the positive periodic association and the negative offset association are no longer distinguishable. This may be one reason why the resting-state alpha literature has sometimes appeared inconsistent. Our findings therefore support the argument that periodic and aperiodic activity should be modelled separately when studying literacy-related individual differences (Donoghue et al., 2020; He, 2014; Turri et al., 2023).

More broadly, this distinction helps connect neural measurement to theory. Periodic alpha is a plausible candidate for indexing rhythmic coordination and timing, whereas aperiodic parameters may reflect different background properties of the neural signal. Treating them separately gives a more precise account of which aspects of the resting EEG spectrum matter for biliteracy development.

### 4.4 Limitations and Future Directions

Several limitations should be acknowledged. First, the cross-sectional nature of our data limits causal inference about developmental directionality. Longitudinal analyses are needed to test whether periodic alpha power prospectively predicts changes in RAN and biliteracy skills and to determine whether aperiodic offset exerts stable negative influences on literacy over time. Second, our alpha measure focused on a posterior ROI selected from the sample-average alpha topography; replication using pre-registered ROIs or independent datasets will strengthen generalizability. Third, although we recorded 3 minutes of eyes-open resting-state EEG, paediatric data quality constraints required artifact rejection; we therefore set a minimum inclusion criterion of fifteen artifact-free 4-s epochs (≥ 60 s) to ensure a minimally stable basis for spectral parameterization while retaining an adequate sample size. Future work should examine the robustness and reliability of periodic alpha estimates across longer recordings and alternative minimum-data thresholds. Finally, the negative associations involving aperiodic offset, although theoretically interpretable, should be considered exploratory and warrant replication in larger, longitudinal samples that can better characterise the functional role of aperiodic activity in literacy development.

## 5. Conclusion

This study shows that posterior periodic alpha power, estimated after separating oscillatory activity from aperiodic background components, is associated with foundational biliteracy skills in Chinese–English bilingual children. The association was evident for word reading and dictation in both languages and was partly mediated by rapid automatized naming, highlighting rapid serial processing efficiency as an important behavioural pathway linking oscillatory neural readiness to reading and spelling outcomes. At the same time, the negative association between aperiodic offset and Chinese literacy, together with the null conventional total-alpha comparison, underscores the value of distinguishing periodic and aperiodic components when studying individual differences in literacy-related neural signals. Together, the findings support he use of spectral parameterization of resting-state EEG as a promising framework for understanding variation in literacy learning.

## Author Contributions

Conceptualization: TKL, SH, UM; Methodology: TKL, SH, UM; Formal Analysis: TKL, SH; Investigation: TKL, SH; Writing – Original Draft Preparation: TKL, SH; Writing – Review & Editing: TKL, SH, KFHL, CM, UM; Visualization: TKL; Supervision: UM; Funding Acquisition: CM, UM.

## Acknowledgements

This research was supported by the Theme-based Research Scheme (T44-410/21-N to U. Maurer, PC, C. McBride, PC) from the Hong Kong Special Administrative Region Research Grants Council.

## Informed Consent Statement

Informed consent was obtained from all subjects involved in the study.

## Data Availability Statement

The de-identified data and analysis scripts supporting the findings of this study are openly available on the Open Science Framework (OSF) at: https://osf.io/e8wxn/overview?view_only=6d5308c4721948e4976d9612d41045be

## Conflicts of Interest

The authors declare no conflicts of interest.

## Abbreviations

The following abbreviations are used in this manuscript:

ADJUST: Automatic EEG artifact Detector based on the Joint Use of Spatial and Temporal features
BCa: Bias-Corrected and Accelerated
CDICT: Chinese Dictation
CDRAN: Chinese Digit Rapid Automatized Naming
CFI: Comparative Fit Index
CI / CIs: Confidence Interval(s)
CWR: Chinese Word Reading
CUDAICA: CUDA-optimized Infomax ICA
EDICT: English Dictation
EEG: Electroencephalography
EEGLAB: EEG Laboratory (software)
EWR: English Word Reading
FID: Family ID
FOOOF: Fitting Oscillations & One-Over-F (algorithm)
Hz: Hertz
ICA: Independent Component Analysis
ICC: Intraclass Correlation Coefficient
PW: Periodic (Alpha) Power
RAN: Rapid Automatized Naming
RMSEA: Root Mean Square Error of Approximation
RS-EEG: Resting-State Electroencephalography
SD: Standard Deviation
SEM: Structural Equation Modelling
SES: Socioeconomic Status
SRMR: Standardized Root Mean Square Residual
TLI: Tucker-Lewis Index
β: Beta (Standardized Regression Coefficient)

---

**Supplementary Table 1:**
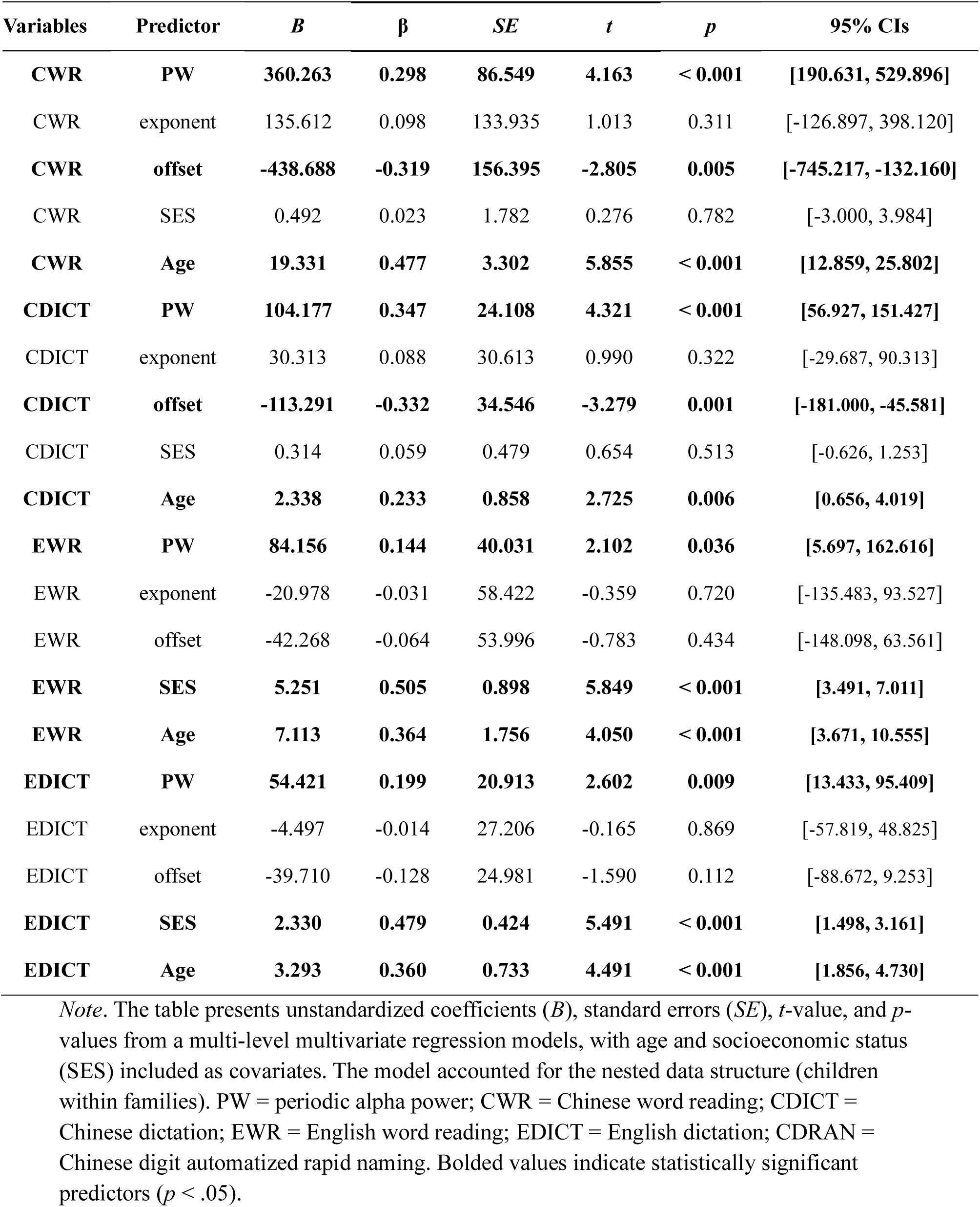
Structural Equation Modelling Predicting Biliteracy Skills from Periodic and Aperiodic Alpha Power.

## Notes

https://osf.io/e8wxn/overview?view_only=6d5308c4721948e4976d9612d41045be

### Competing Interest Statement

The authors have declared no competing interest.

